# Complex evolutionary history of translation Elongation Factor 2 and diphthamide biosynthesis in Archaea and parabasalids

**DOI:** 10.1101/262600

**Authors:** Adrienne B. Narrowe, Anja Spang, Courtney W. Stairs, Eva F. Caceres, Brett J. Baker, Christopher S. Miller, Thijs J. G. Ettema

## Abstract

Diphthamide is a modified histidine residue which is uniquely present in archaeal and eukaryotic elongation factor 2 (EF-2), an essential GTPase responsible for catalyzing the coordinated translocation of tRNA and mRNA through the ribosome. In part due to the role of diphthamide in maintaining translational fidelity, it was previously assumed that diphthamide biosynthesis genes (*dph*) are conserved across all eukaryotes and archaea. Here, comparative analysis of new and existing genomes reveals that some archaea (i.e., members of the Asgard superphylum, *Geoarchaea*, and *Korarchaeota*) and eukaryotes (i.e., parabasalids) lack *dph*. In addition, while EF-2 was thought to exist as a single copy in archaea, many of these *dph*-lacking archaeal genomes encode a second EF-2 paralog missing key-residues required for diphthamide modification and for normal translocase function, perhaps suggesting functional divergence linked to loss of diphthamide biosynthesis. Interestingly, some Heimdallarchaeota previously suggested to be most closely related to the eukaryotic ancestor maintain *dph* genes and a single gene encoding canonical EF-2. Our findings reveal that the ability to produce diphthamide, once thought to be a universal feature in archaea and eukaryotes, has been lost multiple times during evolution, and suggest that anticipated compensatory mechanisms evolved independently.

## INTRODUCTION

Elongation factor 2 (EF-2) is a critical component of the translational machinery that interacts with both the small and large ribosomal subunits. EF-2 functions at the decoding center of the ribosome, where it is necessary for the translocation of messenger RNA and associated tRNAs (Spahn, et al. 2004). Archaeal and eukaryotic EF-2, as well as the homologous bacterial EF-G, are members of the highly conserved translational GTPase protein superfamily (Atkinson 2015). Gene duplications and subsequent neo-functionalizations have been inferred for eukaryotic EF-2 (eEF-2), with the identification of the spliceosome component Snu114 (Fabrizio, et al. 1997), and Ria1, a 60S ribosomal subunit biogenesis factor (Becam, et al. 2001). Bacterial EF-G is involved in both translocation and ribosome recycling and has undergone multiple duplications, including sub-functionalizations separating the translocation and ribosome recycling functions (Suematsu, et al. 2010; Tsuboi, et al. 2009) as well as neo-functionalizations including roles in back-translocation (Qin, et al. 2006), translation termination (Freistroffer, et al. 1997), regulation (Li, et al. 2014) and tetracycline resistance (Donhofer, et al. 2012). However, to date, archaea were thought to encode only a single essential protein within this superfamily, i.e. archaeal EF-2 (aEF-2) (Atkinson 2015).

Unlike bacterial EF-Gs, archaeal and eukaryotic EF-2s contain a post-translationally modified amino acid which is synthesized upon the addition of a 3-amino-3-carboxypropyl (ACP) group to a conserved histidine residue and its subsequent modification to diphthamide by the concerted action of 3 (in archaea) to 7 enzymes (in eukaryotes)(de CrÃ©cy-Lagard, et al. 2012; Schaffrath, et al. 2014). While diphthamide is perhaps best known as the target site of bacterial ADP-ribosylating toxins (Iglewski, et al. 1977; Jorgensen, et al. 2008) and as required for sensitivity to the antifungal sordarin (Botet, et al. 2008), its exact role remains a subject of investigation. Yeast mutants incapable of synthesizing diphthamide have a higher rate of translational frame shifts, suggesting that this residue plays a critical role in reading frame fidelity during translation (Ortiz, et al. 2006). Furthermore, structural studies of eEF-2 using high-resolution Cryo-EM have indicated that diphthamide interacts directly with codon-anticodon bases in the translating ribosome, and facilitates translocation by displacing ribosomal decoding bases (Anger, et al. 2013; Murray, et al. 2016). In addition, diphthamide has been proposed to play a role in the regulation of translation, as it represents a site for reversible endogenous ADP-ribosylation (Schaffrath, et al. 2014), and in the selective translation of certain genes in response to cellular stress (ArgÃ¼elles, et al. 2014). Given its anticipated role at the core of the translational machinery, it is not surprising that, with the sole exception of *Korarchaeum cryptofilum* (de CrÃ©cy-Lagard, et al. 2012; Elkins, et al. 2008), the diphthamide biosynthetic pathway is universally conserved in all archaea and eukaryotes. Indeed, while not strictly essential, loss of diphthamide biosynthesis has been shown to result in growth defects in yeast (Kimata and Kohno 1994; Ortiz, et al. 2006) and some archaea (Blaby, et al. 2010), and is either lethal or causes severe developmental abnormalities in mammals (Liu, et al. 2006; Webb, et al. 2008; Yu, et al. 2014).

In the current study, we explore the evolution and function of EF-2 and of diphthamide biosynthesis genes using genomic data from novel major archaeal lineages that were recently discovered using metagenomics and single-cell genomics approaches (Adam, et al. 2017; Hug, et al. 2016; Spang, et al. 2017). In particular, we report the presence of EF-2 paralogs in many archaeal genomes belonging to the Asgard archaea, *Korarchaeota* and Bathyarchaeota (Evans, et al. 2015; He, et al. 2016; Lazar, et al. 2016; Meng, et al. 2014; Spang, et al. 2015; Zaremba-Niedzwiedzka, et al. 2017) and the unexpected absence of diphthamide biosynthesis genes in several archaea and in parabaslid eukaryotes. Our findings reveal a complex evolutionary history of EF-2 and diphthamide biosynthesis genes, and point to novel mechanisms of translational regulation in several archaeal lineages. Finally, our results are compatible with scenarios in which eukaryotes evolved from an Asgard-related ancestor (Spang, et al. 2015; Zaremba-Niedzwiedzka, et al. 2017) and suggest the presence of a diphthamidated EF-2 in this lineage.

## MATERIALS AND METHODS

### Sampling and sequencing of ABR Loki- and Thorarchaeota

Sampling, DNA extraction, library preparation and sequencing was produced as described in (Zaremba-Niedzwiedzka, et al. 2017). We chose the four deepest samples, at 125 and 175 cm below sea-floor (MM3/PM3 and MM4/PM4 respectively), as they showed highest lokiarchaeal diversity in a maximum likelihood phylogeny of 5 to 15 ribosomal proteins (RP15) encoded on the same contig (Zaremba-Niedzwiedzka, et al. 2017). Adapters and low quality bases were trimmed using Trimmomatic version 0.32 with the following parameters: PE -phred33 ILLUMINACLIP:NexteraPE-PE.fa:2:30:10:1:true LEADING:3 TRAILING:6 SLIDINGWINDOW:4:15 MINLEN:36 (Bolger, et al. 2014).

### Assembly of ABR Loki- and Thorarchaeota

Samples from the same depth were assembled together using IDBA-UD (Peng, et al. 2012) (version 1.1.1-384, –maxk 124 -r <MERGED_READS>) producing four different assemblies (S1:MM1/PM1, S2:MM2/PM2, S3:MM3/PM3, S4:MM4/PM4). Assemblies S3 and S4 were particularly interesting as they showed the highest lokiarchaeal diversity. However, some lokiarchaeal members showed highly fragmented contigs, probably due to the low abundances of these organisms. In an attempt to produce longer contigs we co-assembled those reads coming from Asgard archaea members in the samples MM3, PM3, MM4 and PM4. Asgard archaea reads were identified using Clark (version 1.2.3, -m 0) (Ounit, et al. 2015) and Bowtie2 (version 2.2.4, default parameters) (Langmead and Salzberg 2012) against a customized Asgard archaea database. Classified reads were extracted and co-assembled using SPAdes (version v.3.9.0, –careful) (Bankevich, et al. 2012).

In brief, the Asgard database was composed of Asgard genomes publicly available on February 2017. Clark does not perform well when organisms present in the samples of interest are not highly similar to the ones present in the provided database. To increase the classification sensitivity, we included in our database low-quality Asgard MAGs (with highly fragmented contigs) generated from assemblies S3 and S4, using CONCOCT (Alneberg, et al. 2014). Coverage profiles required by CONCOCT were estimated using kallisto (version 0.43.0, quant –plaintext) (Bray, et al. 2016). All available samples from the same location (MM1, PM1, MM2, PM2, MM3, PM3, MM4, PM4) were used and mapped independently against the assemblies S3 and S4. For each assembly, MAGs were reconstructed using two different minimum contig length thresholds (2000 and 3000 bp). We used the number of containing clusters of ribosomal proteins (ribocontigs) as a proxy to estimate the microbial diversity present in the community. The maximum number of clusters (-c option in CONCOCT) was estimated by calculating approximately 2.5 times the estimated number of species in the sample (Johannes Alneberg, personal communication), resulting into 900 and 600 for S3 and S4, respectively. Potential Asgard archaea bins were identified based on the presence of ribocontigs classified as Asgard archaea and were included in the database.

### Binning of ABR Loki- and Thorarchaeota

Several binning tools with different settings were run independently: CONCOCT_2000: version 0.4.0, –read_length 200 and minimum contig length of 2000. CONCOCT_3000: version 0.4.0, -read_length 200 and minimum contig length of 3000. In both cases, coverage files were created mapping all 8 samples against the co-assembly using kallisto. MaxBin2: version 2.2.1, -min_contig_length 2000 -markerset 40 –plotmarker (Wu, et al. 2016). The 8 samples were mapped against the co-assembly using Bowtie2. Coverage was estimated using the getabund.pl script provided. MyCC_4mer: 4mer -t 2000 (Lin and Liao 2016). MyCC_56mer: 56mer -t 2000. Both coverage profiles were obtained as the authors described in their manual.

The results of those 5 binning methods were combined into a consensus: contigs were assigned to bins if they had been classified as the same organism by at least 3 out of 5 methods. The resulting bins were manually inspected and cleaned further using mmgenome (Albertsen, et al. 2013). Completeness and redundancy was computed using CheckM (Parks, et al. 2015).

### Sampling and sequencing of OWC Thorarchaeota

Eight soil samples were collected from the Old Woman Creek (OWC) National Estaurine Research Reserve and DNA was extracted as described previously (Narrowe, et al. 2017). Library preparation and five lanes of Illumina HiSeq 2x125 bp sequencing followed standard operating procedures at the US DOE Joint Genome Institute (GOLD study ID Gs0114821). Sample M3-C4-D3 had replicate extraction, library preparation, and two lanes of sequencing performed, and reads were combined before downstream analysis. For 3 additional samples (M3-C4-D4, O3-C3-D3, O3-C3-D4) one lane of sequencing was performed. For the other 4 samples (M3-C5-D1, M3-C5-D2, M3-C5-D3, M3-C5-D4) DNA was sheared to 300bp with a Covaris S220, metagenomic sequencing libraries were prepared using the Nugen Ovation Ultralow Prep kit, and all four samples were multiplexed on one lane of Illumina HiSeq 2x125 sequencing at the University of Colorado Denver Anschutz Medical Campus Genomics and Microarray Core.

### Assembly and binning of OWC Thorarchaeota

For initial assembly of the 5 full-lane sequencing runs, adapter removal, read filtering and trimming were completed using BBDuk (sourceforge.net/projects/bbmap) ktrim=r, minlen=40, minlenfraction=0.6, mink=11 tbo, tpe k=23, hdist=1 hdist2=1 ftm=5, maq=8, maxns=1, minlen=40, minlenfraction=0.6, k=27, hdist=1, trimq=12, qtrim=rl. Filtered reads were assembled using megahit (Li, et al. 2015) version 1.0.6 with –k-list 23,43,63,83,103,123.

The individual metagenome from the O3-C4-D3 sample was binned using Emergent Self-Organizing Maps (ESOM)(Dick, et al. 2009) of tetranucleotide frequency (5kb contigs, 3kb windows). BLAST hits of predicted proteins identified a Thorarchaeota population bin. All scaffolds containing a window in this bin were used as a mapping reference and reads from the 9 OWC libraries were mapped to this bin using bbsplit with default parameters (sourceforge.net/projects/bbmap). The mapped reads were reassembled using SPAdes version 3.9.0 with –careful -k 21,33,55,77,95,105,115,125 (Bankevich, et al. 2012). Finally, the reads which were input to the reassembly were mapped to the assembled scaffolds using Bowtie 2 (Langmead and Salzberg 2012) to generate a coverage profile which was used to manual identify bins using Anvi’o (Eren, et al. 2015). Proteins were predicted using prodigal (Hyatt, et al. 2010) and searched against UniRef90 release 11-2016 (Suzek, et al. 2015), with the taxonomy of best blast hits used to validate contigs as probable Thorarchaeota. Contigs having no top hit to the publicly available Thorarchaeota genomes were manually examined and removed if they could be assigned to another genome bin in the larger metagenomic assembly. Genome completeness and contamination was estimated using CheckM (Parks, et al. 2015).

### Identification of diphthamide biosynthesis genes and EF-2 homologs in eukaryotes and archaea

The EGGNOG members dataset (available at http://eggnogdb.embl.de/#/app/downloads) was surveyed for sequences corresponding to the following clusters of orthologous groups (COG): EF-2, COG0480; DPH1/DPH2, COG1736; DPH3, COG5216; DPH4, COG0484; DPH5, COG1798; DPH6, COG2102; and DPH7, ENOG4111MMJ. For genomes not represented in EGGNOG, we manually inspected publicly available genomes as indicated by ‘orthology assignment source’ (Supplementary File S1). Similarly, an in-house arCOG dataset, modeled after the publicly available arCOGs from Makarova et al. (Makarova, et al. 2015),was queried for the corresponding COG distribution in relevant archaeal genomes. Finally, aEF-2 and aEF-2p genes in Thorarchaeota OWC Bin 2,3 and 5 were identified using HMMER: version 3.1b2, hmmsearch –cut-tc (Eddy 2011) against PFAM models PF00679 (EF-G_C) and PF03764 (EFG_IV). Conserved synteny surrounding the Thorarchaoeta aEF-2p gene was used to further search for partial aEF-2p genes. In addition, all contigs with matching HMM hits to *dph2* and *dph5* in the full OWC assembly were manually examined for potential Thorarchaeal *dph* genes; none were identified.

### Phylogenetic analyses

Elongation factor 2: EF-2 and EF-2 paralogs of Asgard archaea, Koarchaeota and Bathyarchaeota were aligned with a representative set of archaeal, bacterial EF-2 and eukaryotic EF-2, EFL1 and snRNP homologs using mafft-linsi (Katoh and Standley 2013). Subsequently, poorly aligned ends were removed manually before the alignments were trimmed with trimAl 5% (Capella-Gutierrez, et al. 2009), yielding 871 aligned amino acid positions. Maximum likelihood analyses were performed using IQ-tree using the mixture model LG+C60+R4+F, which was selected among the C-series models based on its Bayesian information criterion score by the built-in model test implemented in IQ-tree. Branch supports were assessed using ultrafast bootstrap approximation as well as with single branch test (-alrt option).

Diphthamide biosynthesis proteins Dph1/Dph2 (IPR016435; arCOG04112) and Dph5 (IPR004551; arCOG04161): Both Dph1 and Dph2 as well as Dph5 homologs of a representative set of eukaryotes were aligned with archaeal Dph1/2 and Dph5 homologs, respectively. Several DPANN genomes contain two genes encoding the CTD and NTD of Dph1/2 (Fig. 1, Supplementary File S1) such that Dph1/2 homologs of these organisms had to be concatenated prior to aligning Dph1/2 sequences. Alignments were performed using mafft-linsi and trimmed with BMGE (Criscuolo and Gribaldo 2010) using the blossum 30 matrix and setting the entropy to 0.55. This resulted in final alignments of 170 (Dph1/2) and 221 (Dph5). Maximum likelihood analyses were performed using IQ-tree (Nguyen, et al. 2015) with the mixture models resulting in the lowest BIC: LG+C50+R+F (Dph1/2) and LG+C60+R+F (Dph5), respectively. Branch supports were assessed using ultrafast bootstrap approximation (Hoang, et al. 2018) as well as with the single branch test (-alrt flag).

**Figure 1.**
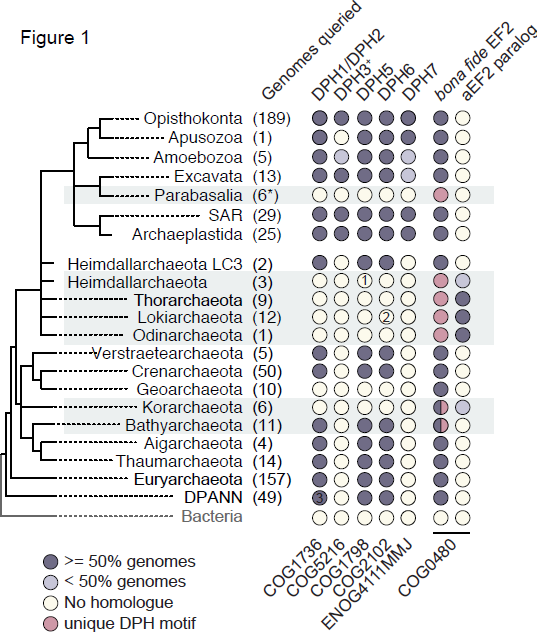
Diphthamide biosynthesis genes are conserved across most eukaryotic and archaeal lineages. Eukaryotic and archaeal orthologues of diphthamide biosynthesis (DPH) genes were retrieved from the publicly available EGGNOG and an in-house archaeal orthologues (arCOG) datasets. Complete list of genomes surveyed can be found in Supplementary File S1 including reduced genomes from nucleomorphs (not shown on figure). Total number of genomes surveyed are shown next to each group. Since Dph4 is a member of the large DNAJ-containing protein family, we could not unequivocally identify this protein based on orthology alone and is therefore excluded from the figure. ^+^No arCOG available for DPH3. *All eukaryotic genomes are complete except five deeply-sequenced transcriptomes from Parabasalia; dark and light grey circles indicate whether homologues were detected in more or less than 50% of the genomes surveyed respectively; yellow circles indicate the absence of a detectable homologue; pink circles indicate lack of conservation of the diphthamide modification motif; half-circles indicate the presence of multiple copies of EF-2 with and without the conserved diphthamide modification motif. 1 - Homologue detected in the original assembly (ABR_125(Zaremba-Niedzwiedzka, et al. 2017)) but not in the reassembly (ABR16 genome); a closer inspection of the contig revealed that it is chimeric and will thus be removed from the final bin; 2 - Homologue detected in only one Lokiarchaeota assembly (AB_15); 3 - Several DPANN genomes contain two proteins that encode the CTD and NTD of Dph1/2, respectively.

Concatenated ribosomal proteins: A phylogenetic tree of co-localized ribosomal proteins was performed using the rp15 pipeline as described previously (Zaremba-Niedzwiedzka, et al. 2017). In brief, archaeal ribosomal proteins encoded in the r-protein gene cluster (requiring a minimum of 11 ribosomal proteins) were aligned with mafft-linsi, trimmed with trimAl using the -gappyout option, concatenated and subjected to maximum likelihood analyses using IQ-tree with the LG+C60+R4+F model chosen based on best BIC score as described above. Branch supports were assessed using ultrafast bootstrap approximation as well as with the single branch test (-alrt option) in IQTREE.

### Structural modeling of EF-2 homologs

Structural models of a/eEF-2 genes and paralogs were generated using the i-Tasser standalone package version 5.1 (Yang, et al. 2015), and visualized and analyzed using UCSF Chimera version 1.11.12 (Pettersen, et al. 2004). The best structural hits to the PDB for each sequence’s top-scoring model were identified using COFACTOR (Roy, et al. 2012). The *Drosophila melanogaster eEF-2* structure in complex with the ribosome (PDB:4V6W) was used as a structural reference to which all models were superimposed (aligned) using Chimera’s MatchMaker.

### Loop motif logos of EF-2 homologs

e/aEF-2 and paralog sequences which were used to generate the EF-2 tree were clustered at 90% amino acid identity using CD-HIT: version 4.6, -c 0.9 -n 5 (Fu, et al. 2012) and the sequence alignment was filtered to retain only cluster centroids. The conserved loop sequences were extracted from the filtered EF-2 alignment using Jalview version 2.10.1 (Waterhouse, et al. 2009), verified by cross-referencing to the structural models, and sequence logos generated on cluster centroids only using WebLogo: version 2.8.2 (weblogo.berkeley.edu) (Crooks, et al. 2004).

### Accession Numbers

Taxonomy and accession numbers for all genes analyzed in this study are listed in Supplementary File S1.

## RESULTS

### Most Asgard archaea, *Korarchaeota* and *Geoarchaea* as well as parabasalids lack diphthamide synthesis genes

It was previously assumed that EF-2 of all eukaryotes and Archaea was uniquely characterized by the presence of diphthamide. To examine if this assumption is still valid when taking into account recently sequenced genomes, we surveyed 337 archaeal and 168 eukaryotic genomes (File S1) for each of the three known archaeal (de CrÃ©cy-Lagard, et al. 2012) and seven eukaryotic (Su, et al. 2012a; Su, et al. 2012b; Uthman, et al. 2013) *dph* genes. While most archaeal genomes encode clear *dph* homologues, we failed to detect the diphthamide biosynthesis genes in a large diversity of metagenome-assembled genomes (MAGs) of uncultured archaea, including newly assembled MAGs analyzed for this study (Fig. 1, Supplementary Fig. S1, Supplementary File S1). In particular, our analyses showed that, as reported for *K. cryptophilum* (de CrÃ©cy-Lagard, et al. 2012; Elkins, et al. 2008), all *Korarchaeota* and *Geoarchaea* as well as nearly all members of the Asgard archaea lack the conserved archaeal diphthamide biosynthesis genes *dph1/2*, *dph5* and *dph6*. As an exception, Asgard archaea related to the Heimdallarchaeote LC3 clade were found to encode the complete archaeal diphthamide biosynthetic pathway (Fig. 1). Genes coding for Dph5 and Dph6 could not be detected in two Bathyarchaeota draft genomes (RBG_13_46_16b and SG8_32_3). However, it is unclear whether these two genomes are in the process of losing *dph* biosynthesis genes or whether the absence of *dph5* and *dph6* genes is due to the incompleteness of these draft genomes. We also surveyed 168 eukaryotic genomes and high-quality transcriptomes, including those lineages that have undergone drastic genome reduction, such as microsporidians (Corradi, et al. 2010), diplomonads (Morrison, et al. 2007), and degenerate nuclei (i.e., nucleomorphs) of secondary plastids in cryptophytes (Lane, et al. 2007) (Supplementary File S1) for *dph* gene homologs. We detected *dph* homologues in all eukaryotic genomes and transcriptomes except for parabasalid protists, including animal pathogens such as *Trichomonas vaginalis, Tritrichomonas foetus* and *Dientamoeba fragilis* (Supplementary File S1). Unless these archaea and parabasalids possess alternative, yet undiscovered diphthamide biosynthesis pathways, these findings suggest that their cognate EF-2 lacks the modified diphthamide residue. As a peculiarity, while the Dph1/2 protein is encoded by a single fusion gene in seemingly all archaea, we found that in several members of the DPANN archaea (Castelle, et al. 2015; Rinke, et al. 2013) this protein is encoded by two genes that separately code for the N- and C-terminal domains. To our knowledge, this is the first systematic report of the widespread absence of diphthamide biosynthesis in diverse eukaryotes and archaea.

### Various archaeal genomes that lack diphthamide biosynthesis genes encode an EF-2 paralog

To shed light into the implications of the potential lack of diphthamide in members of the Asgard archaea and *Korarchaeota*, we performed detailed analyses of eukaryotic and archaeal EF-2 homologs (Fig. 1). First, we found that the draft genomes of most Asgard archaea, some *Korarchaeota* (Kor 1 and 3), and a few Bathyarchaeota encode two distantly related EF-2 paralogs. In contrast, the genomes of *K. cyptophilum* and two novel marine *Korarchaeota* (Kor 2 and 4) and Heimdallarchaeota LC2 and LC3 as well as *Geoarchaea* do not encode an EF-2 paralog. Given that the Heimdallarchaeota LC2 genome was estimated to be only 70-79 % complete (Zaremba-Niedzwiedzka, et al. 2017), and based on phylogenetic analyses (see below), we consider it possible that this genome might encode an as-yet unassembled aEF-2 paralog. The presence of paralogous aEF-2 in most Asgard archaea and some *Korarchaeota* genomes corresponds with the absence of diphthamide synthesis genes (Fig. 1 and 2). Yet, even though the genomes of *K. cryptophilum*, Kor 2, Kor 4, and *Geoarchaea* as well as of Heimdallarchaeote LC2 lack *dph* genes, they do not encode an EF-2 paralog. In all other archaeal genomes, including that of Heimdallarchaeote LC3, the absence of an EF-2 paralog correlates with the presence of *dph* genes.

### Archaea with two EF-2 family proteins encode only one *bona fide* EF-2

We next addressed whether residues and structural motifs shown to be necessary for canonical translocation were conserved in the various EF-2 and EF-2 paralogs. Domain IV of EF-2, representing the anticodon mimicry domain, is critical for facilitating concerted translocation of tRNA and mRNA (Ortiz, et al. 2006; Rodnina, et al. 1997). This domain includes three loops that extend out from the body of EF-2 and interact with the decoding center of the ribosome. The first of these three loops (HxDxxHRG) (canonical residue positions are numbered according to sequence associated with *D. melanogaster* structural model PDB 4V6W (Anger, et al. 2013)) contains the site of the diphthamide modified histidine, H701, and is highly conserved across archaea and eukaryotes (Ortiz, et al. 2006; Zhang, et al. 2008). High conservation is also seen in a second adjacent loop (SPHKHN) in the a/eEF-2 domain IV (S581-N586), which contains a lysine residue (K584) that interacts directly with the tRNA at the decoding center, and is itself positioned by a stacking interaction between P582 and H585 (Murray, et al. 2016). The third loop appears to stabilize the diphthamide loop, partially via a salt-bridge formed between a nearby glutamate residue (E660) and R702 in the diphthamide loop (Anger, et al. 2013). Both of these residues are highly conserved among archaea and eukaryotes.

Our analyses reveal that the sequence motifs in these loops are also strictly conserved among the EF-2 family proteins of the Heimdallarchaeote LC3 lineage, *Geoarchaea*, as well as in those *Korarchaeota* and Bathyarchaeota that lack an EF-2 paralog (Fig. 3, Supplementary Fig. S2a). Notably, this conservation is seen irrespective of the presence or absence of *dph* genes in those genomes. However, most *bona fide* EF-2 of parabasalids (which lack *dph* genes), possesses a glycine to asparagine mutation at residue 703 (Fig. 3, Supplementary Fig. S2b, Supplementary Fig. S3a), which may compensate for the lack of the diphthamide residue by contributing an amide group (Fig. 3, Supplementary Fig. S3b).

In contrast, in those Asgard archaea and *Korarchaeota* (Kor 1/3 clade) that encode two EF-2 family proteins, even within the bona fide EF-2 copy, these domain IV motifs show reduced conservation. In the diphthamide loop, R702 is universally replaced by a threonine residue. In 21 of 22 aEF-2 proteins, there is a correlated mutation of E660 to either arginine or lysine (Supplementary Fig. S4). Structural homology modeling suggested that these correlated mutations likely prevent unfavorable electrostatic interactions between domain IV loops, and maintain stabilization of the diphthamide loop (Supplementary Fig. S4). While G703 is conserved in most EF-2s of archaea, all Lokiarchaeota (except Lokiarchaeota CR_4), encode either a serine or a glutamine at this site (Fig. 3, Supplementary Fig. S2a). Furthermore, analysis of the second loop (S581-N586) revealed additional crucial mutations in the EF-2 of these archaea; notably, K584 is not conserved (Fig. 3, Supplementary Fig. S2a). Despite these modifications which correlate with the presence of an EF-2 paralog in these archaea, there is still evidence for strong selection pressure maintaining many of the key conserved residues in these domain IV motifs, including H701, the target site of diphthamide modification (Fig. 3, Supplementary Fig. S2a).

In contrast, our analyses of the multiple sequence alignment and structural models suggest that the paralogous EF-2 (aEF-2p) proteins encoded by these archaea lack conservation in the stabilizing second loop (SPHKHN) as well as the first diphthamide loop (HxDxxHRG), including H701 (Fig. 3). Based on predicted fold conservation in domains I and II, and the overall conservation of the five sequence motifs (G1-G5) characterizing GTPase superfamily proteins (Atkinson 2015), aEF-2p likely maintains GTPase activity (Supplementary Fig. S5). However, given the apparent lack of conservation in key domain IV loops, it is unlikely that aEF-2p proteins can serve as functional translocases in protein translation.

### EF-2 homologs of archaea experienced complex evolutionary history

To resolve the evolutionary history of EF-2, we performed phylogenetic analyses of archaeal EF-2 (aEF-2) and aEF-2p, bacterial EF-G and eukaryotic EF-2 family proteins, i.e. EF-2, Ria1 (or Elongation factor like, EFL1) and Snu114 (or U5 small nuclear ribonucleoprotein, snRNP/ U5-116kD) (Fig. 2) (Atkinson 2015). First, our analyses revealed that sequences from all non-LC3 Asgard archaea and the Kor-1 and -3 marine *Korarchaeota* formed two distinct clades, one of which contains canonical aEF-2 proteins (as defined by conservation of the domain IV loop known to interact with the ribosomal decoding center during translocation) while the other cluster comprises aEF-2p (Fig. 2). However, the phylogenetic placement of these protein clades relative to each other and within the phylogenetic backbone is not fully resolved due to lack of statistical support. This might be caused by modified (accelerated) evolutionary rates that appear to characterize the evolution of aEF-2 and aEF-2p in lineages that encode a paralog, as indicated by increased relative branch lengths for both the aEF-2 and aEF-2p clades (Fig. 2, Supplementary Files S2 and S3).

**Figure 2.**
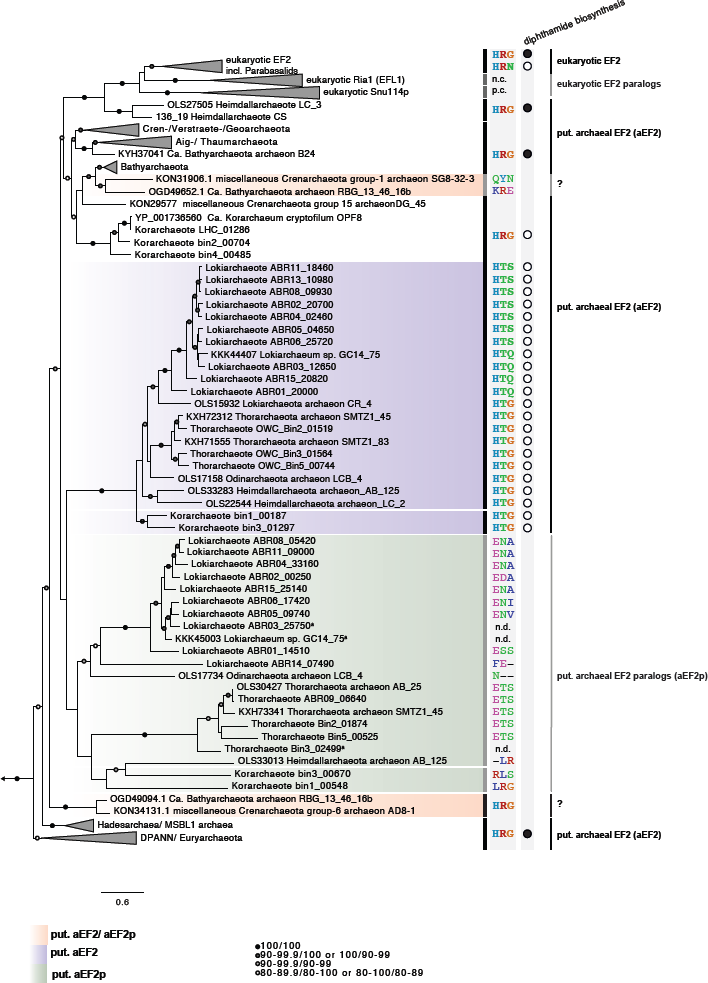
The evolution of archaeal EF-2 family proteins. Phylogenetic tree of EF-2 family proteins based on maximum likelihood analyses of 871 aligned positions using IQ-tree. EF-2 of Bathyarchaeota grouping in an unexpected position or representing potential aEF-2p are shaded in orange. aEF-2 of Kor- and Asgard archaea are shaded in purple, while their aEF-2p are shaded in green. Highlighted amino acids show the conservation of key residues and black/white circles reveal the presence/absence of *dph* biosynthesis genes in the respective organisms/MAGs. Branch support values are based on ultrafast bootstrap approximation as well as single branch tests, respectively and are represented by differentially colored circles as detailed in the figure panel. Whenever branch support values were below 80 for any of the two methods, values have been removed and branches cannot be considered significantly supported. Scale bar indicates the number of substitutions per site. Abr.; snRNP: U5 small nuclear ribonucleoprotein EFL1: elongation factor-like GTPase; n.c.: not conserved; p.c.: partially conserved; n.d.: not determined.

**Figure 3.**
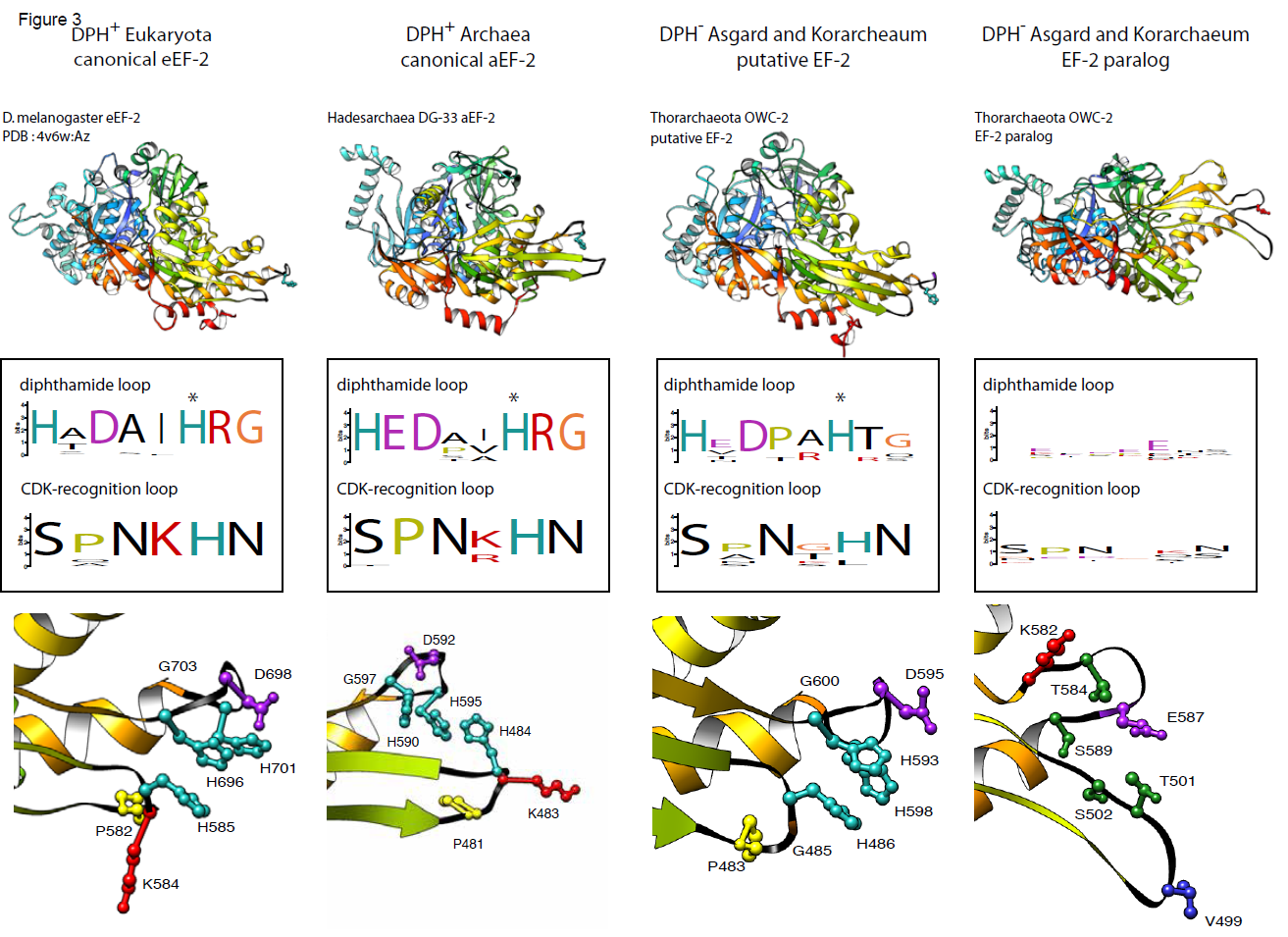
Predicted structure of Asgard archaea EF-2 and EF-2 paralogs. Structural modeling of representative EF-2 genes and paralogs compared to eukaryotic EF-2 structure shows conservation of overall EF-2 structure regardless of diphthamide synthesis capacity (top). The overall fold of two loops located at the tip of domain IV is conserved, but otherwise highly conserved sequence motifs in these loops are not conserved in DPH^-^ Asgard archaea and Korarchaea or in EF-2 paralogs (middle). Bottom panels show a close-up of the key residues from the motifs, highlighting that these residues are those positioned at the tip of the domain IV loops crucial for interaction with the decoding site in canonical EF-2 structures. Histidine residue that is the site of dph modification is starred.

Secondly, bathyarchaeal EF-2 homologs were also found to form two separate clades. One of these clades is placed within the TACK superphylum, and includes both canonical bathyarchaeal EF-2s as well as potential paralogs (i.e., RBG_13_46_16b and SG8-32-3). In contrast, the second clade is only comprised of two sequences (i.e., RBG_13_46_16b and AD8-1), and is placed as a sister group of all TACK, Asgard and eukaryotic EF-2 homologs (Fig. 2). In spite of this deep placement in the phylogenetic analyses, the second clade is comprised of the canonical EF-2 homologs of Bathyarchaeota genomes RBG_13_46_16b and AD8-1, based on analysis of key domain IV residues. Currently, only the most complete of the latter two draft genomes, RBG_13_46_16b, contains an aEF-2 paralog. Therefore, the current data is insufficient to resolve the puzzling pattern of EF-2 evolution in the Bathyarchaeota phylum.

Finally, in our analysis, eEF-2, Ria1 and Snu114 were found to form a highly supported monophyletic group that emerged as a sister group to the aEF-2 proteins encoded by the genomes comprising the Heimdallarchaeote LC3 clade (LC3 and B3). Close inspection of the EF-2 sequence alignment revealed that eukaryotic and LC3 EF-2 homologs share common indels to the exclusion of all other archaeal EF-2 family protein sequences (Supplementary Fig. S6, Supplementary Fig. S7). Notably, these highly conserved indels were found to be encoded by the genomic bins of two distantly related members of the Heimdallarchaeota LC3 lineage, which were independently assembled and binned from geographically distinct metagenomes (Spang, et al. 2015; Zaremba-Niedzwiedzka, et al. 2017). This refutes recently raised claims stating that these indels in Heimdallarchaeote LC3 may be the results of contamination from eukaryotes (Da Cunha, et al. 2017) while supporting the sister-relationship of eukaryotes and Asgard archaea (Eme, et al. 2017; Spang, et al.; Spang, et al. 2015; Zaremba-Niedzwiedzka, et al. 2017). In addition, despite the low sequence identity of 39%, the high-confidence modeled structure of Heimdallarchaeote LC3 EF-2 was highly similar to *Drosophila melanogaster* eEF-2 (RMSD (root-mean-square deviation) 1.3Ã across all 796 residues to *D. melanogaster* structural model PDB 4V6W (Anger, et al. 2013); Supplementary File S1). By comparison, the Heimdallarchaeaote AB-125 model aligns less confidently to the Drosophila EF-2 structure (RMSD 16.4Ã). The observed phylogenetic topology and the presence of the full complement of *dph* biosynthesis genes in LC3 genomes (Figs. 1 and 2), support an evolutionary scenario in which Heimdallarchaeote LC3 and eukaryotes share a common ancestry with EF-2 being vertically inherited from this archaeal ancestor.

## DISCUSSION

The use of metagenomic approaches has led to an expansion of genomic data from a large diversity of previously unknown archaeal and bacterial lineages and has changed our perception of the tree of life, microbial metabolic diversity and evolution, as well as the origin of eukaryotes (Brown, et al. 2015; Castelle, et al. 2015; Hug, et al. 2016; Parks, et al. 2017; Spang, et al. 2015; Zaremba-Niedzwiedzka, et al. 2017). Since most of what is known about archaeal informational processing machineries is based on a few model organisms, we aimed to use the expansion of genomic data to investigate key elements of the translational machinery - EF-2 and diphthamidylation - across the tree of life.

Our analyses of archaeal EF-2 family proteins and the distribution of diphthamide biosynthesis genes have revealed unusual features of the core translation machinery in several archaeal lineages. These findings negate two long-held assumptions regarding the archaeal and eukaryotic translation machineries, with both functional and evolutionary implications. First, we show that diphthamide modification is not universally conserved across Archaea and eukaryotes. Second, we demonstrate that, much like Bacteria and eukaryotes (Atkinson 2015), the archaeal EF-2 protein family has undergone several gene duplication events, presumably coupled to functional differentiation of EF-2 paralogs, throughout archaeal evolution.

The evolution of archaeal diphthamide biosynthesis and EF-2 is especially intriguing in the context of eukaryogenesis. Recent findings based on comparative genomics indicate that eukaryotes evolved from a symbiosis between an alphaproteobacterium with an archaeal host that shares a most recent common ancestor with extant members of the Asgard archaea, possibly a Heimdallarchaeota-related lineage (Spang, et al. 2015; Zaremba-Niedzwiedzka, et al. 2017). Our study adds additional data to support this scenario by revealing close sequence and predicted structural similarity of canonical EF-2 proteins of the Heimdallarchaeote LC3 lineage and eukaryotic EF-2 proteins, including shared indels. Furthermore, phylogenetic analyses of EF-2 family proteins reveals that EF-2 of the Heimdallarchaeote LC3 lineage forms a monophyletic group with EF-2 family proteins of eukaryotes, and therefore suggests that the archaeal ancestor of eukaryotes was equipped with an EF-2 protein similar to the homologs found in this lineage. The subsequent evolution of the eukaryotic EF-2 family appears to have included at least two ancient duplication events leading to Ria1 and Snu114. Importantly, the presence of characteristic eukaryotic indels in EF-2 of all members of the Heimdallarchaeote LC3 lineage further strengthens this hypothesis and underlines that concerns raised about the quality of these genomic bins (Da Cunha, et al. 2017) are unjustified (Spang, et al.).

In addition, the LC3 clade also represents the sole group within the Asgard archaea that is characterized by the presence of the full complement of archaeal diphthamide biosynthesis pathway genes. However, while phylogenetic analyses of Dph1/2 show weak support for a sister-relationship between Heimdallarchaeota and eukaryotes, eukaryotic Dph5 appears to be most closely related to homologs of Woesearchaeaota (Supplementary Fig. S8, Supplementary File S3), an archaeal lineage belonging to the proposed DPANN superphylum (Castelle, et al. 2015; Rinke, et al. 2013; Williams, et al. 2017), comprising various additional lineages with putative symbiotic and/or parasitic members (reviewed in Spang et al. (Spang, et al. 2017)). Notably, a previous study has also revealed an affiliation of some eukaryotic tRNA synthetases with DPANN archaea (Furukawa, et al. 2017). Given that several DPANN lineages infect or closely associate with other archaeal lineages, they may exchange genes with their hosts frequently, as was shown for *Nanoarchaeum equitans* and its crenarchaeal host *Ignicoccus hospitalis* (Podar, et al. 2008). Following a similar reasoning, the archaeal ancestor of eukaryotes (i.e. a relative of the Asgard archaea) may have acquired genes (e.g. *dph5*) from an ancestral DPANN/Woesearchaeota symbiont. However, prospective analyses and generation of genomic data from additional members of the Asgard and DPANN archaea are necessary to test this hypothesis and to clarify the evolutionary history of the origin of diphthamide biosynthesis genes in eukaryotes.

Furthermore, our findings have practical implications for studies that involve phylogenetic and metagenomic analyses. Previously, EF-2 has been widely used as a phylogenetic marker, in both single-gene (Baldauf, et al. 1996; Elkins, et al. 2008; Hashimoto and Hasegawa 1996; Iwabe, et al. 1989), and multiple-gene alignments of universal single copy genes [(Guy, et al. 2014; Raymann, et al. 2015; Williams, et al. 2012), and others] to assess the relationships between Archaea, Bacteria and eukaryotes. However, the presence of paralogs of EF-2 in various Archaea and eukaryotes suggest that EF-2 should be excluded from such datasets. In addition, EF-2, Dph1/2, and Dph5 are part of single-copy marker gene sets regularly used to estimate genome completeness and purity of archaeal metagenomic bins (Parks, et al. 2015; Wu and Scott 2012). The presence of duplicated aEF-2 gene families, the absence of *dph* genes in most Asgard archaea, *Geoarchaea* and *Korarchaeota*, and the presence of two split genes for Dph1/2 in DPANN makes these genes unsuited as marker genes, and should hence be excluded from marker gene sets used to assess genome completeness.

The observed absence of *dph* biosynthesis genes in various Archaea as well as parabasalids is surprising given that diphthamide was previously thought to be a conserved feature across Archaea and eukaryotes (Schaffrath, et al. 2014), and critical for ensuring translational fidelity (Ortiz, et al. 2006). While we currently cannot rule out the possibility that *dph*-lacking archaea and parabasalids perform the multi-step process of diphthamidylation using a set of yet-unknown enzymes, future proteomics studies will be needed to conclusively rule out the presence of diphthamide in these taxa. Yet, it is more likely that these groups have evolved a different mechanism or mechanisms to fulfill the proposed roles of diphthamide in translation.

Many of the *dph*-lacking archaeal genomes encode two paralogs of the aEF-2 gene. Despite the apparent absence of diphthamide, our sequence and structural modeling analyses imply that these dipthamide-deficient aEF-2 proteins are likely under strong selective pressure to maintain translocase function. In contrast, analyses of the aEF-2p suggest that, while this paralog is a member of the translational GTPase superfamily, aEF-2p is unlikely to function in the same manner as canonical aEF-2. In fact, the complete lack of sequence conservation in aEF-2p key domain IV loop residues indicates that these paralogs are not likely to act as translocases (Fig. 3, Supplementary Fig. S2a) (Ortiz, et al. 2006; Rodnina, et al. 1997) and instead perform alternative roles. For instance, it seems possible that aEF-2p may compensate for the absence of diphthamide in at least some *dph*-lacking lineages. However, other functions for aEF-2p such as error-correcting back-translocation or ribosome recycling also seem possible, given the observed sub- and neo-functionalizations seen in eukaryotic and bacterial EF-2/EF-G paralogs (Qin, et al. 2006; Tsuboi, et al. 2009). Alternatively, given proposed regulation of translation via ADP-ribosylation of diphthamide (Schaffrath, et al. 2014) and a role of diphthamide in responding to oxidative stress (ArgÃ¼elles, et al. 2014; ArgÃ¼elles, et al. 2013), aEF-2p could perform another, yet unknown role in translation regulation.

Currently, the consequences for the absence of *dph* biosynthesis genes in parabasalids and in several Archaea remain unclear. Future studies could gain insight into such questions by studying translation in the genetically tractable parabasalid *Trichomonas vaginalis,* whose cell biology and metabolism has been extensively studied. In addition, acquisition of additional sequencing data or enrichment cultures from members of the Asgard superphylum, *Korarchaeota*, and other novel archaeal lineages will lead to a better understanding of the evolution and function of EF-2 family proteins, and the absence of *dph* biosynthesis genes.

## ACKNOWLEDGEMENTS

We thank Jordan Angle, Kay Stefanik, Rebecca Daly, and Kelly Wrighton for assistance with sampling of OWC sediments, and Felix Homa for computational support. Sequencing of OWC metagenomes was conducted in part by the U.S. Department of Energy Joint Genome Institute, a DOE Office of Science User Facility that is supported by the Office of Science of the U.S. Department of Energy under Contract No. DE-AC02-05CH11231. Sequencing of Aarhus bay metagenomes was performed by the National Genomics Infrastructure sequencing platforms at the Science for Life Laboratory at Uppsala University, a national infrastructure supported by the Swedish Research Council (VR-RFI) and the Knut and Alice Wallenberg Foundation. We thank the Uppsala Multidisciplinary Center for Advanced Computational Science (UPPMAX) at Uppsala University and the Swedish National Infrastructure for Computing (SNIC) at the PDC Center for High-Performance Computing for providing computational resources. This work was supported by grants of the European Research Council (ERC Starting grant 310039-PUZZLE_CELL), the Swedish Foundation for Strategic Research (SSF-FFL5) and the Swedish Research Council (VR grant 2015-04959) to T.J.G.E̤ C.W.S. is supported by a European Molecular Biology Organisation long-term fellowship (ALTF-997-2015) and the Natural Sciences and Engineering Research Council of Canada postdoctoral research fellowship (PDF-487174-2016).

## SUPPLEMENTARY INFORMATION

**Supplementary Figures.pdf** - Contains all supplementary figures referenced in the text.

**Supplementary File S1 - Sheet 1: Distribution of diphthamide biosynthesis genes in archaea.** Archaeal homologues and their corresponding accession numbers for each Dph gene are shown as retrieved from an in-house archaeal COG (arCOG) dataset. Total counts for each archaeal group of interest are shown. **Sheet 2: Distribution of diphthamide biosynthesis genes in eukaryotes.** Eukaryotic homologues and their corresponding accession numbers for each Dph gene are shown as retrieved from EGGNOG or manual inspection. Total counts for each eukaryotic supergroup are indicated in different colours. When appropriate, nucleomorph- or nucleus-encoded sequences are indicated. **Sheet 3: Structural modeling results including scoring of top structural model predicted by i-Tasser and best structural hit to that model from PDB.**

**Supplementary File S2 - Trimmed alignment of EF-2 homologs that was used for phylogenetic analyses.** The alignment was generated using mafft-LINSi (Katoh K, Standley DM. Mol Biol Evol. 30:772-80, 2013, doi:10.1093/molbev/mst010) and subjected to trimming with trimAL (5%) (Capella-Gutierrez S, Silla-Martinez JM, Gabaldon T. Bioinformatics 25:1972-3, 2009, doi:10.1093/bioinformatics/btp348) after the manual removal of poorly aligned ends. Please refer to methods section for more details.

**Supplementary File S3a - Newick file of concatenated ribosomal proteins phylogeny.** Please refer to figure legend of Fig. S1 for more details.

**Supplementary File S3b - Newick file of EF-2 phylogeny presented in Figure 2.** Please refer to figure legend of Fig. 2 for more details.

**Supplementary File S3c - Newick file of Dph1/2 phylogeny.** Please refer to figure legend of Fig. S8a for more details.

**Supplementary File S3d - Newick file of Dph5 phylogeny.** Please refer to figure legend of Fig. S8b for more details.

**Supplementary Figures for:**

**Supplementary Fig. S1.**
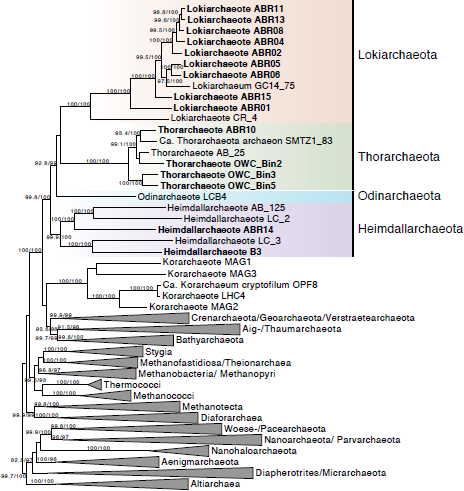
Phylogenetic analysis of at least 11 out of 15 concatenated archaeal ribosomal proteins (2416 AA) based on maximum likelihood analyses performed with IQ-tree. The diverse metagenome-assembled genomes (MAGs) belonging to Asgard archaea and included in our analyses are shaded in colors according to phylum. MAGs that were not part of the initial description of the Asgard superphylum (Zaremba-Niedzwiedzka K, Caceres EF, Saw JH, Backstrom D, Juzokaite L, Vancaester E, Seitz KW, Anantharaman K, Starnawski P, Kjeldsen KU, Stott MB, Nunoura T, Banfield JF, Schramm A, Baker BJ, Spang A, Ettema TJ. Nature 541:353-358, 2017, doi:10.1038/nature21031) are shown in boldface. Naming of the respective archaeal groups based on a recent suggestion by Adam et al. (ISMEJ 11:2407-2425,2017,doi:10.1038/ismej.2017.122). Branch support values are based on ultrafast bootstrap approximation as well as single branch tests, respectively. Scale bar indicates the number of substitutions per site.

**Supplementary Fig. S2.**
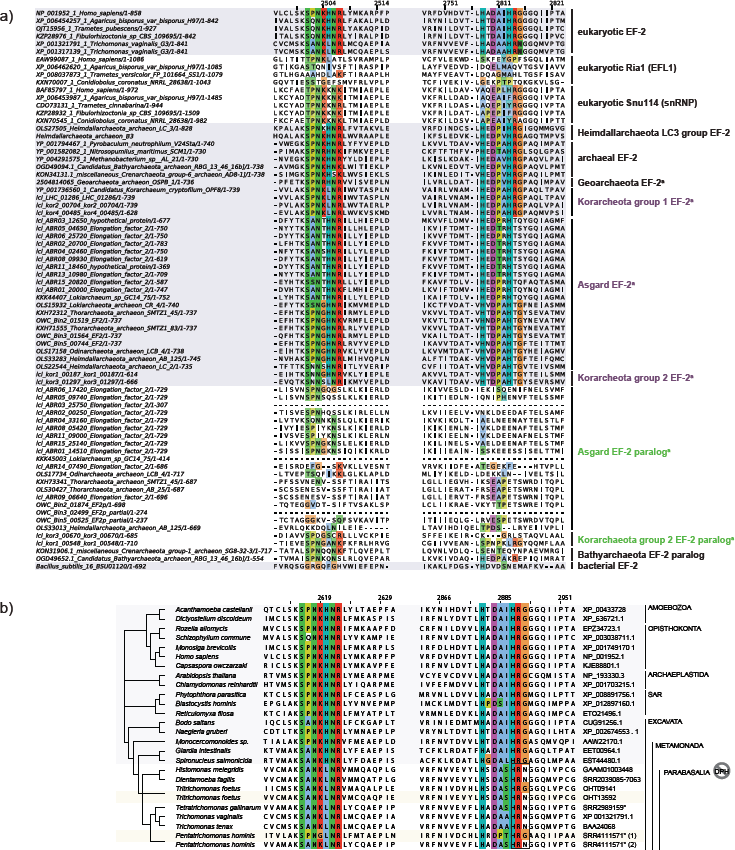
Multiple sequence alignment of archaeal and eukaryotic EF-2 and EF-2 paralogs showing domain IV sequence motifs. (a) Multiple sequence alignment of a selected set of EF-2 from representative organisms, showing domain IV sequence motifs as in Figure 3. *Bona fide* EF-2 homologues are shaded in grey. Organisms lacking diphthamide biosynthesis genes are indicated with ‘a’. (b) Diphthamide modification motifs are not conserved in parabasalid EF-2. EF-2 sequences were mined and aligned from representative genomes or transcriptomes from each of the major lineages of eukaryotes and diphthamide-interacting residues are colored. Here, we show a representative subset of eukaryotes, all surveyed genomes can be found in File S1. Eukaryotic relationships are shown with a schematic cladogram. *Bona fide* diphthamidylated EF-2 sequences are shaded in purple. Boxed region indicates the region that is not conserved in most parabasalids. Parabasalid EF-2 paralogs with unsubstituted diphthamide modification motifs are shaded in yellow. All parabasalids to not encoded diphthamide biosynthesis genes as indicated with the ‘no DPH’ icon. SAR, Stramenopilia, Alveolata, Rhizaria; DPH, diphthamide biosynthesis genes. The *Pentatrichomonas hominis* and *Tetratrichomonas gallinarum* sequences were retrieved by assembling the sequencing projects available at the indicated SRA accession numbers. Sequences and assembly are available upon request.

**Supplementary Fig. S3.**
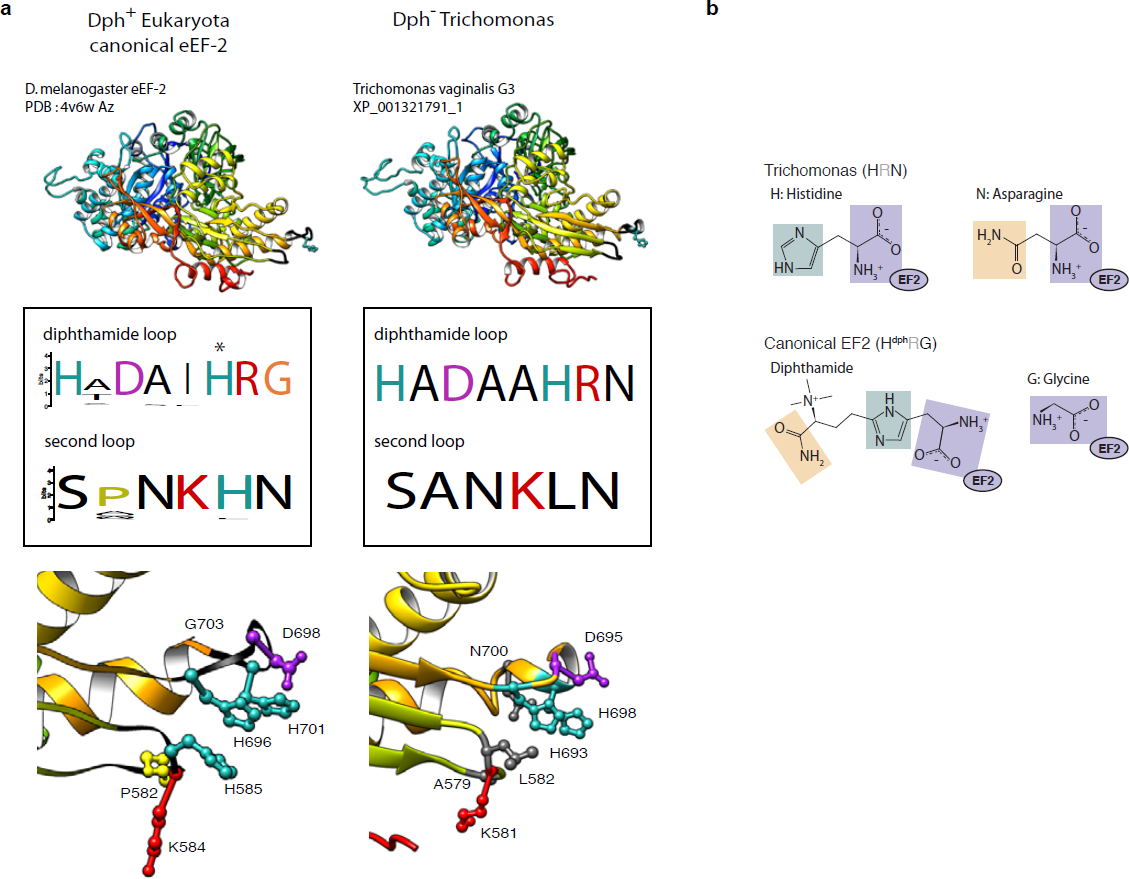
EF-2 gene from Dph-lacking *Trichomonas vaginalis* shown aligned to *D. melanogaster* eEF-2 structure. (a) Panels are as in Figure 3. *T. vaginalis* EF-2 fits closely to *D. melanogaster* structure (RMSD of 1.589 Ã
across all 830 residues). While overall structure is maintained, certain key residues in domain IV loops are not conserved. (b) Structure of the three last amino acids comprising the diphthamide loop in EF-2 of *T. vaginalis* compared to canonical eukaryotic EF-2. The amino acids comprising the DRG motif of canonical EF2 (with D referring to diphthamide) have a backbone highly similar to the HRN motif of *T. vaginalis* (with the histidine being not modified to diphthamide). The mutation of the canonical G to N, which provides an amide group, may compensate for the lack of the modification of the histidine.

**Supplementary Fig. S4.**
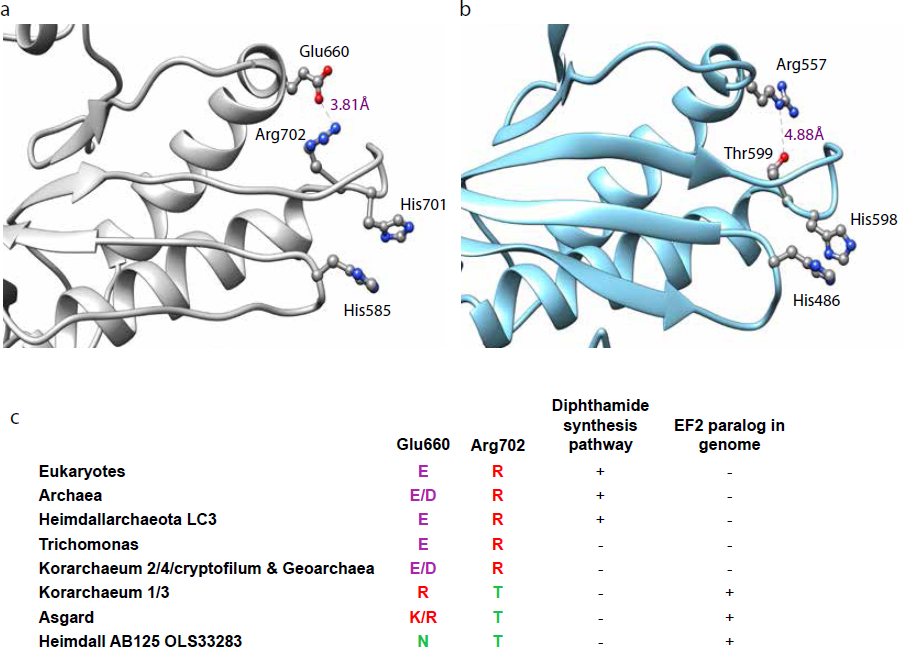
A universally conserved EF-2 domain IV salt bridge is replaced by conserved correlated mutations in EF-2p containing genomes. (a) EF-2 from the *D. melanogaster* EF2 cryo-EM structure (Anger AM, Armache JP, Berninghausen O, Habeck M, Subklewe M, Wilson DN, Beckmann R. Nature 497:80-5, 2013, doi:10.1038/nature12104) shows that Glu660 and Arg702, which are universally conserved in all archaeal and eukaryotic genomes lacking aEF-2p (Figure 3), form a salt bridge that stabilizes the diphthamide-containing loop of domain IV. (b) Representative modeled EF-2 structure of Thorarchaeota OWC Bin 2, with correlated mutations to Arg557 and Thr599 highlighted. (c) Thr599 is conserved in all EF-2p-containing genomes, and the correlated mutation at the Arg557 position is almost always positive or polar.

**Supplementary Fig. S5:**
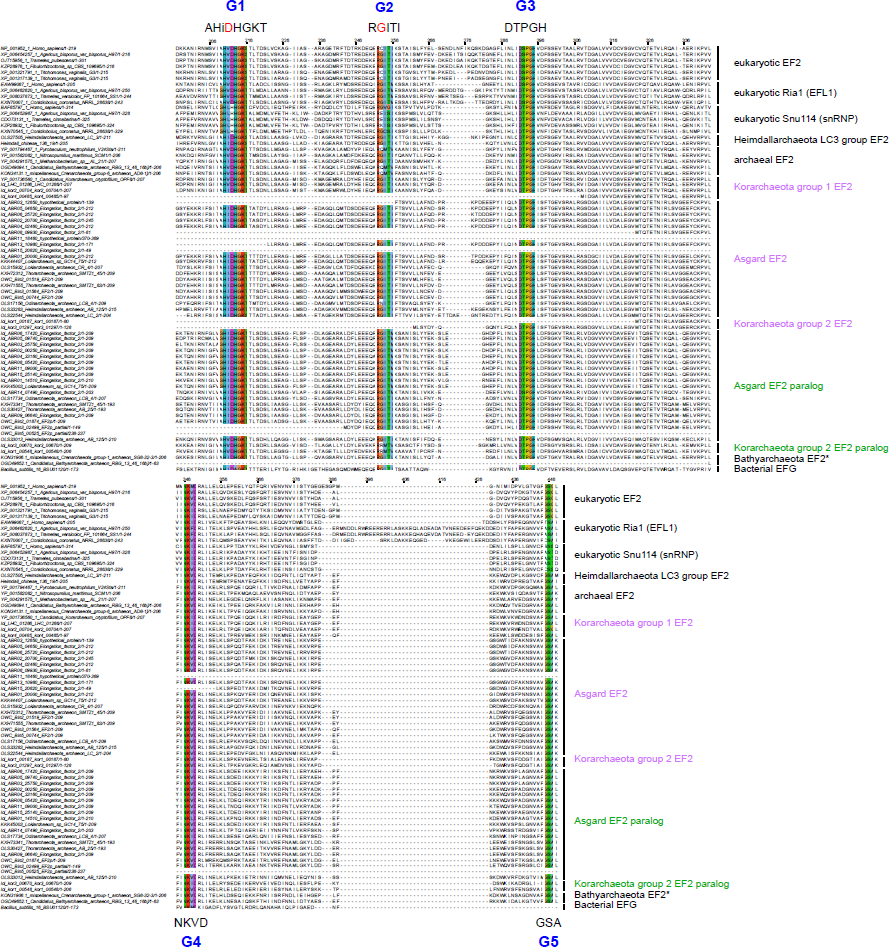
Multiple sequence alignment showing conservation of GTP binding region motifs in Asgard aEF-2 and EF-2 paralogs. Multiple sequence alignment of eEF-2, eEF-2 paralogs, aEF-2, aEF-2 paralogs and bacterial EF-G. Conserved GTP binding motifs G1 - G5 are shown in color in the alignment. Archaeal 60% consensus motif sequences as identified by Atkinson [BMC Genomics 16:78, 2015, doi:10.1186/s12864-015-1289-7] are shown outside the alignment and residues associated with cation binding are shown in red.

**Supplementary Fig. S6:**
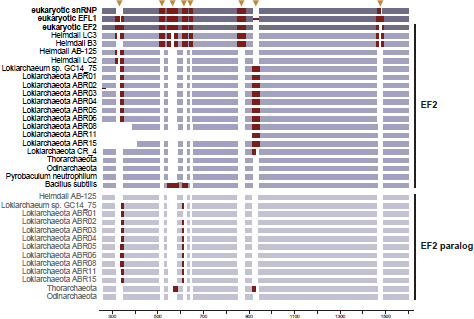
Schematic view of occurrence of indels in archaeal, eukaryotic and bacterial EF-2 and EF-2 paralogs. The cartoon is based on an alignment of EF-2 and EF-2 paralogs from a selected set of representative organisms, mainly comprising Asgard archaea and Eukaryotes. Canonical EF-2 sequences are represented by purple and EF-2 paralogs by light purple bars. Potential indels are shaded by red bars and indel positions are highlighted with orange triangles.

**Supplementary Fig. S7:**
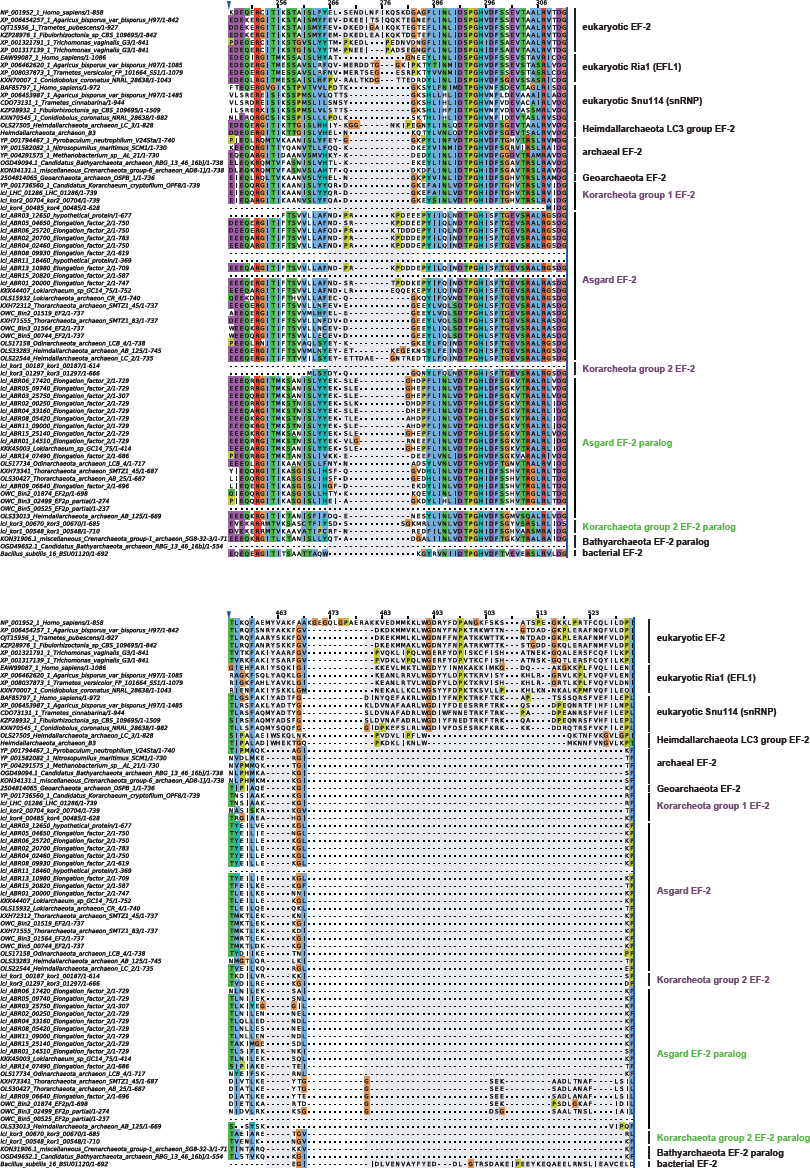
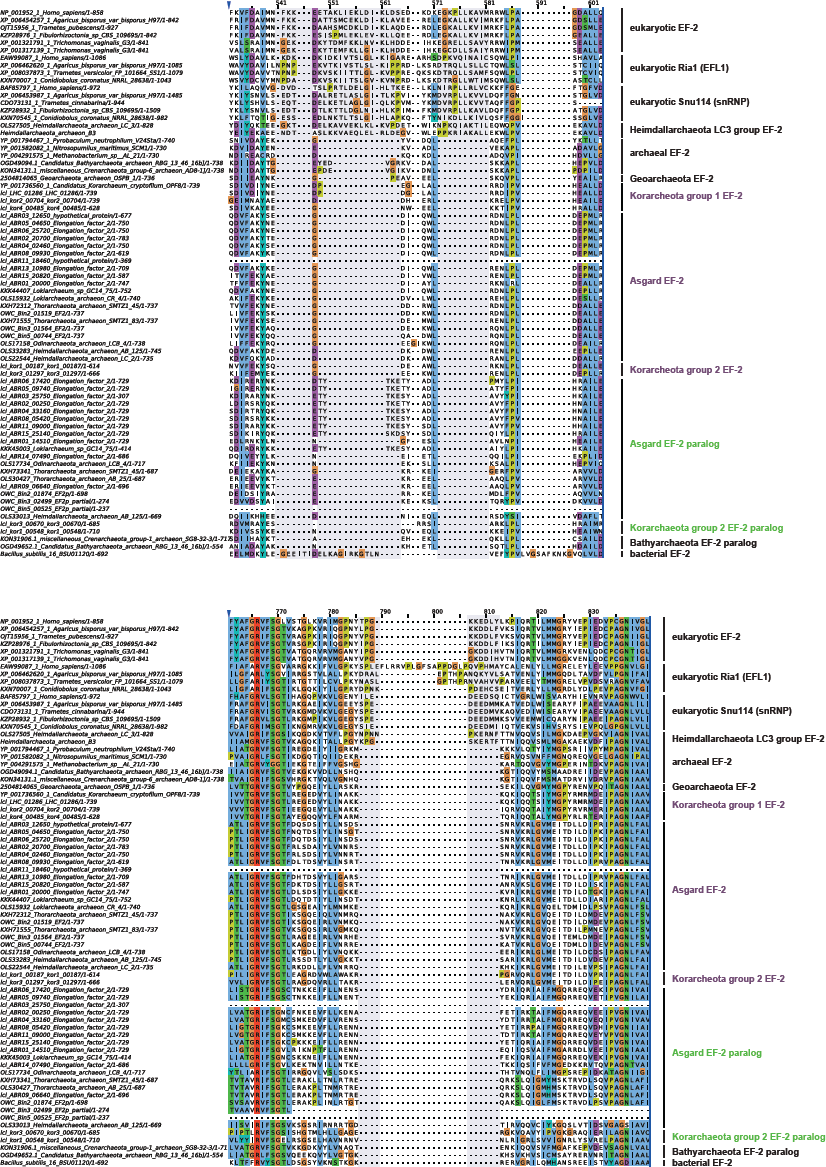
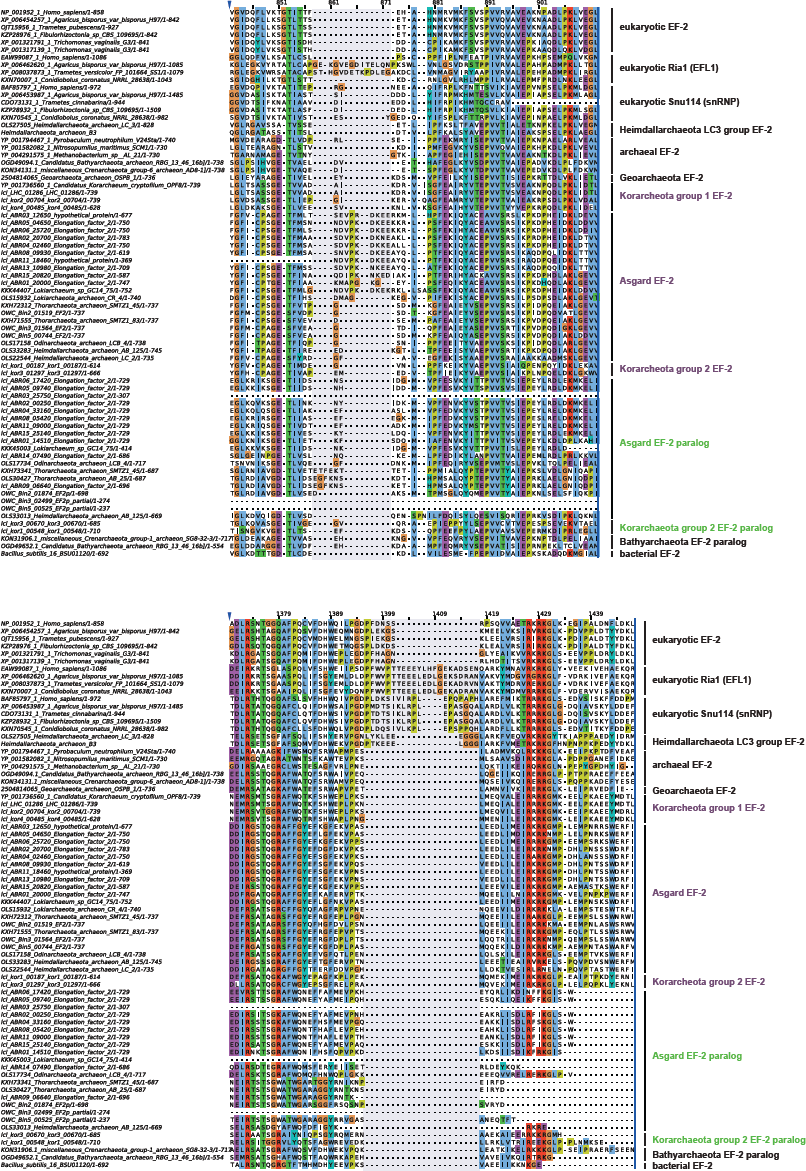
Multiple sequence alignment of occurrence of indels in archaeal, eukaryotic, and bacterial EF-2 and EF-2 paralogs. Selected characteristic indel regions derived from the multiple sequence alignment of a representative set of EF-2 homologs, which provided the basis for Fig. S6. Indel positions are shaded in light purple.

**Supplementary Fig. S8:**
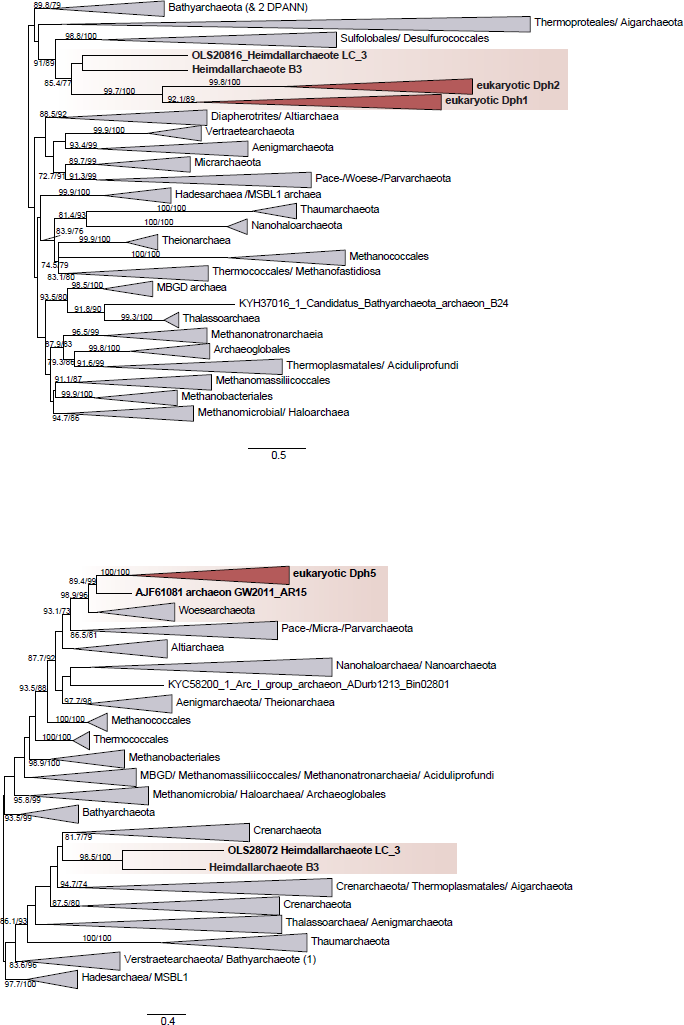
Maximum likelihood phylogenetic analyses of Dph1/2 (a) and Dph5 (b) performed using IQ-tree. Eukaryotic homologs were collapsed and are represented by dark red triangles, while homologs of Woesearchaeota and Heimdallarchaeota are shaded in light red. Both phylogenetic trees are unrooted. Values on branches refer to support values based on ultrafast bootstrap approximation as well as single branch tests. Whenever any of the two support values were lower than 70, bootstraps were removed. Scale bar indicates the number of substitutions per site.

## REFERENCES

Adam PS, Borrel G, Brochier-Armanet C, Gribaldo S 2017. The growing tree of Archaea: new perspectives on their diversity, evolution and ecology. ISME J 11: 2407–2425. doi: 10.1038/ismej.2017.122

Albertsen M, et al. 2013. Genome sequences of rare, uncultured bacteria obtained by differential coverage binning of multiple metagenomes. Nat Biotechnol 31: 533–538. doi: 10.1038/nbt.2579

Alneberg J, et al. 2014. Binning metagenomic contigs by coverage and composition. Nat Methods 11: 1144–1146. doi: 10.1038/nmeth.3103

Anger AM, et al. 2013. Structures of the human and Drosophila 80S ribosome. Nature 497: 80–85. doi: 10.1038/nature12104

ArgÃ¼elles S, Camandola S, Cutler RG, Ayala A, Mattson MP 2014. Elongation factor 2 diphthamide is critical for translation of two IRES-dependent protein targets, XIAP and FGF2, under oxidative stress conditions. Free Radic Biol Med 67: 131–138. doi: 10.1016/j.freeradbiomed.2013.10.015

ArgÃ¼elles S, et al. 2013. Molecular control of the amount, subcellular location, and activity state of translation elongation factor 2 in neurons experiencing stress. Free Radic Biol Med 61: 61–71. doi: 10.1016/j.freeradbiomed.2013.03.016

Atkinson GC 2015. The evolutionary and functional diversity of classical and lesser-known cytoplasmic and organellar translational GTPases across the tree of life. BMC Genomics 16: 78. doi: 10.1186/s12864-015-1289-7

Baldauf SL, Palmer JD, Doolittle WF 1996. The root of the universal tree and the origin of eukaryotes based on elongation factor phylogeny. Proc Natl Acad Sci U S A 93: 7749–7754.

Bankevich A, et al. 2012. SPAdes: a new genome assembly algorithm and its applications to single-cell sequencing. J Comput Biol 19: 455–477. doi: 10.1089/cmb.2012.0021

Becam AM, Nasr F, Racki WJ, Zagulski M, Herbert CJ 2001. Ria1p (Ynl163c), a protein similar to elongation factors 2, is involved in the biogenesis of the 60S subunit of the ribosome in Saccharomyces cerevisiae. Mol Genet Genomics 266: 454–462. doi: 10.1007/s004380100548

Blaby IK, et al. 2010. Towards a systems approach in the genetic analysis of archaea: Accelerating mutant construction and phenotypic analysis in Haloferax volcanii. Archaea 2010: 426239. doi: 10.1155/2010/426239

Bolger AM, Lohse M, Usadel B 2014. Trimmomatic: a flexible trimmer for Illumina sequence data. Bioinformatics 30: 2114–2120. doi: 10.1093/bioinformatics/btu170

Botet J, Rodriguez-Mateos M, Ballesta JP, Revuelta JL, Remacha M 2008. A chemical genomic screen in Saccharomyces cerevisiae reveals a role for diphthamidation of translation elongation factor 2 in inhibition of protein synthesis by sordarin. Antimicrob Agents Chemother 52: 1623–1629. doi: 10.1128/AAC.01603-07

Bray NL, Pimentel H, Melsted P, Pachter L 2016. Near-optimal probabilistic RNA-seq quantification. Nat Biotechnol 34: 525–527. doi: 10.1038/nbt.3519

Brown CT, et al. 2015. Unusual biology across a group comprising more than 15% of domain Bacteria. Nature 523: 208–211. doi: 10.1038/nature14486

Capella-Gutierrez S, Silla-Martinez JM, Gabaldon T 2009. trimAl: a tool for automated alignment trimming in large-scale phylogenetic analyses. Bioinformatics 25: 1972–1973. doi: 10.1093/bioinformatics/btp348

Castelle CJ, et al. 2015. Genomic expansion of domain archaea highlights roles for organisms from new phyla in anaerobic carbon cycling. Curr Biol 25: 690–701. doi: 10.1016/j.cub.2015.01.014

Corradi N, Pombert JF, Farinelli L, Didier ES, Keeling PJ 2010. The complete sequence of the smallest known nuclear genome from the microsporidian Encephalitozoon intestinalis. Nat Commun 1: 77. doi: 10.1038/ncomms1082

Criscuolo A, Gribaldo S 2010. BMGE (Block Mapping and Gathering with Entropy): a new software for selection of phylogenetic informative regions from multiple sequence alignments. BMC Evol Biol 10: 210. doi: 10.1186/1471-2148-10-210

Crooks GE, Hon G, Chandonia JM, Brenner SE 2004. WebLogo: a sequence logo generator. Genome Res 14: 1188–1190. doi: 10.1101/gr.849004

Da Cunha V, Gaia M, Gadelle D, Nasir A, Forterre P 2017. Lokiarchaea are close relatives of Euryarchaeota, not bridging the gap between prokaryotes and eukaryotes. PLoS Genet 13: e1006810. doi: 10.1371/journal.pgen.1006810

de CrÃ©cy-Lagard V, Forouhar F, Brochier-Armanet C, Tong L, Hunt JF 2012. Comparative genomic analysis of the DUF71/COG2102 family predicts roles in diphthamide biosynthesis and B12 salvage. Biol Direct 7: 32. doi: 10.1186/1745-6150-7-32

Dick GJ, et al. 2009. Community-wide analysis of microbial genome sequence signatures. Genome Biol 10: R85. doi: 10.1186/gb-2009-10-8-r85

Donhofer A, et al. 2012. Structural basis for TetM-mediated tetracycline resistance. Proc Natl Acad Sci U S A 109: 16900–16905. doi: 10.1073/pnas.1208037109

Eddy SR 2011. Accelerated Profile HMM Searches. PLoS Comput Biol 7: e1002195. doi: 10.1371/journal.pcbi.1002195

Elkins JG, et al. 2008. A korarchaeal genome reveals insights into the evolution of the Archaea. Proceedings of the National Academy of Sciences 105: 8102–8107. doi: 10.1073/pnas.0801980105

Eme L, Spang A, Lombard J, Stairs CW, Ettema TJG 2017. Archaea and the origin of eukaryotes. Nature Reviews Microbiology 15: 711. doi: 10.1038/nrmicro.2017.133

Eren AM, et al. 2015. Anvi’o: an advanced analysis and visualization platform for ‘omics data. PeerJ 3: e1319. doi: 10.7717/peerj.1319

Evans PN, et al. 2015. Methane metabolism in the archaeal phylum Bathyarchaeota revealed by genome-centric metagenomics. Science 350: 434–438. doi: 10.1126/science.aac7745

Fabrizio P, Laggerbauer B, Lauber J, Lane WS, Luhrmann R 1997. An evolutionarily conserved U5 snRNP-specific protein is a GTP-binding factor closely related to the ribosomal translocase EF-2. EMBO J 16: 4092–4106. doi: 10.1093/emboj/16.13.4092

Freistroffer DV, Pavlov MY, MacDougall J, Buckingham RH, Ehrenberg M 1997. Release factor RF3 in E.coli accelerates the dissociation of release factors RF1 and RF2 from the ribosome in a GTP-dependent manner. EMBO J 16: 4126–4133. doi: 10.1093/emboj/16.13.4126

Fu L, Niu B, Zhu Z, Wu S, Li W 2012. CD-HIT: accelerated for clustering the next-generation sequencing data. Bioinformatics 28: 3150–3152. doi: 10.1093/bioinformatics/bts565

Furukawa R, Nakagawa M, Kuroyanagi T, Yokobori SI, Yamagishi A 2017. Quest for Ancestors of Eukaryal Cells Based on Phylogenetic Analyses of Aminoacyl-tRNA Synthetases. J Mol Evol 84: 51–66. doi: 10.1007/s00239-016-9768-2

Guy L, Saw JH, Ettema TJ 2014. The archaeal legacy of eukaryotes: a phylogenomic perspective. Cold Spring Harb Perspect Biol 6: a016022. doi: 10.1101/cshperspect.a016022

Hashimoto T, Hasegawa M 1996. Origin and early evolution of eukaryotes inferred from the amino acid sequences of translation elongation factors 1alpha/Tu and 2/G. Adv Biophys 32: 73–120.

He Y, et al. 2016. Genomic and enzymatic evidence for acetogenesis among multiple lineages of the archaeal phylum Bathyarchaeota widespread in marine sediments. Nature Microbiology 1: 16035. doi: 10.1038/nmicrobiol.2016.35

Hoang DT, Chernomor O, von Haeseler A, Minh BQ, Vinh LS 2018. UFBoot2: Improving the Ultrafast Bootstrap Approximation. Mol Biol Evol 35: 518–522. doi: 10.1093/molbev/msx281

Hug LA, et al. 2016. A new view of the tree of life. Nature Microbiology 1. doi: Artn 16048 10.1038/Nmicrobiol.2016.48

Hyatt D, et al. 2010. Prodigal: prokaryotic gene recognition and translation initiation site identification. BMC Bioinformatics 11: 119. doi: 10.1186/1471-2105-11-119

Iglewski BH, Liu PV, Kabat D 1977. Mechanism of action of Pseudomonas aeruginosa exotoxin Aiadenosine diphosphate-ribosylation of mammalian elongation factor 2 in vitro and in vivo. Infect Immun 15: 138–144.

Iwabe N, Kuma K, Hasegawa M, Osawa S, Miyata T 1989. Evolutionary relationship of archaebacteria, eubacteria, and eukaryotes inferred from phylogenetic trees of duplicated genes. Proc Natl Acad Sci U S A 86: 9355–9359.

Jorgensen R, et al. 2008. Cholix toxin, a novel ADP-ribosylating factor from Vibrio cholerae. J Biol Chem 283: 10671–10678. doi: 10.1074/jbc.M710008200

Katoh K, Standley DM 2013. MAFFT multiple sequence alignment software version 7: improvements in performance and usability. Mol Biol Evol 30: 772–780. doi: 10.1093/molbev/mst010

Kimata Y, Kohno K 1994. Elongation factor 2 mutants deficient in diphthamide formation show temperature-sensitive cell growth. J Biol Chem 269: 13497–13501.

Lane CE, et al. 2007. Nucleomorph genome of Hemiselmis andersenii reveals complete intron loss and compaction as a driver of protein structure and function. Proc Natl Acad Sci U S A 104: 19908–19913. doi: 10.1073/pnas.0707419104

Langmead B, Salzberg SL 2012. Fast gapped-read alignment with Bowtie 2. Nat Methods 9: 357–359. doi: 10.1038/nmeth.1923

Lazar CS, et al. 2016. Genomic evidence for distinct carbon substrate preferences and ecological niches of Bathyarchaeota in estuarine sediments. Environ Microbiol 18: 1200–1211. doi: 10.1111/1462-2920.13142

Li D, Liu CM, Luo R, Sadakane K, Lam TW 2015. MEGAHIT: an ultra-fast single-node solution for large and complex metagenomics assembly via succinct de Bruijn graph. Bioinformatics 31: 1674–1676. doi: 10.1093/bioinformatics/btv033

Li L, et al. 2014. Ribosomal elongation factor 4 promotes cell death associated with lethal stress. MBio 5: e01708. doi: 10.1128/mBio.01708-14

Lin HH, Liao YC 2016. Accurate binning of metagenomic contigs via automated clustering sequences using information of genomic signatures and marker genes. Sci Rep 6: 24175. doi: 10.1038/srep24175

Liu S, et al. 2006. Dph3, a small protein required for diphthamide biosynthesis, is essential in mouse development. Mol Cell Biol 26: 3835–3841. doi: 10.1128/MCB.26.10.3835-3841.2006

Makarova KS, Wolf YI, Koonin EV 2015. Archaeal Clusters of Orthologous Genes (arCOGs): An Update and Application for Analysis of Shared Features between Thermococcales, Methanococcales, and Methanobacteriales. Life (Basel) 5: 818–840. doi: 10.3390/life5010818

Meng J, et al. 2014. Genetic and functional properties of uncultivated MCG archaea assessed by metagenome and gene expression analyses. ISME J 8: 650–659. doi: 10.1038/ismej.2013.174

Morrison HG, et al. 2007. Genomic minimalism in the early diverging intestinal parasite Giardia lamblia. Science 317: 1921–1926. doi: 10.1126/science.1143837

Murray J, et al. 2016. Structural characterization of ribosome recruitment and translocation by type IV IRES. Elife 5. doi: 10.7554/eLife.13567

Narrowe AB, et al. 2017. High-resolution sequencing reveals unexplored archaeal diversity in freshwater wetland soils. Environ Microbiol 19: 2192–2209. doi: 10.1111/1462-2920.13703

Nguyen LT, Schmidt HA, von Haeseler A, Minh BQ 2015. IQ-TREE: a fast and effective stochastic algorithm for estimating maximum-likelihood phylogenies. Mol Biol Evol 32: 268–274. doi: 10.1093/molbev/msu300

Ortiz PA, Ulloque R, Kihara GK, Zheng H, Kinzy TG 2006. Translation elongation factor 2 anticodon mimicry domain mutants affect fidelity and diphtheria toxin resistance. J Biol Chem 281: 32639–32648. doi: 10.1074/jbc.M607076200

Ounit R, Wanamaker S, Close TJ, Lonardi S 2015. CLARK: fast and accurate classification of metagenomic and genomic sequences using discriminative k-mers. BMC Genomics 16: 236. doi: 10.1186/s12864-015-1419-2

Parks DH, Imelfort M, Skennerton CT, Hugenholtz P, Tyson GW 2015. CheckM: assessing the quality of microbial genomes recovered from isolates, single cells, and metagenomes. Genome Res 25: 1043–1055. doi: 10.1101/gr.186072.114

Parks DH, et al. 2017. Recovery of nearly 8,000 metagenome-assembled genomes substantially expands the tree of life. Nature Microbiology 2: 1533–1542. doi: 10.1038/s41564-017-0012-7

Peng Y, Leung HC, Yiu SM, Chin FY 2012. IDBA-UD: a de novo assembler for single-cell and metagenomic sequencing data with highly uneven depth. Bioinformatics 28: 1420–1428. doi: 10.1093/bioinformatics/bts174

Pettersen EF, et al. 2004. UCSF Chimera—a visualization system for exploratory research and analysis. J Comput Chem 25: 1605–1612. doi: 10.1002/jcc.20084

Podar M, et al. 2008. A genomic analysis of the archaeal system Ignicoccus hospitalis-Nanoarchaeum equitans. Genome Biol 9: R158. doi: 10.1186/gb-2008-9-11-r158

Qin Y, et al. 2006. The highly conserved LepA is a ribosomal elongation factor that back-translocates the ribosome. Cell 127: 721–733. doi: 10.1016/j.cell.2006.09.037

Raymann K, Brochier-Armanet C, Gribaldo S 2015. The two-domain tree of life is linked to a new root for the Archaea. Proc Natl Acad Sci U S A 112: 6670–6675. doi: 10.1073/pnas.1420858112

Rinke C, et al. 2013. Insights into the phylogeny and coding potential of microbial dark matter. Nature 499: 431–437. doi: 10.1038/nature12352

Rodnina MV, Savelsbergh A, Katunin VI, Wintermeyer W 1997. Hydrolysis of GTP by elongation factor G drives tRNA movement on the ribosome. Nature 385: 37–41. doi: 10.1038/385037a0

Roy A, Yang J, Zhang Y 2012. COFACTOR: an accurate comparative algorithm for structure-based protein function annotation. Nucleic Acids Res 40: W471–477. doi: 10.1093/nar/gks372

Schaffrath R, Abdel-Fattah W, Klassen R, Stark MJ 2014. The diphthamide modification pathway from Saccharomyces cerevisiae—revisited. Mol Microbiol 94: 1213–1226. doi: 10.1111/mmi.12845

Spahn CM, et al. 2004. Domain movements of elongation factor eEF2 and the eukaryotic 80S ribosome facilitate tRNA translocation. EMBO J 23: 1008–1019. doi: 10.1038/sj.emboj.7600102

Spang A, Caceres EF, Ettema TJG 2017. Genomic exploration of the diversity, ecology, and evolution of the archaeal domain of life. Science 357. doi: 10.1126/science.aaf3883

Spang A, et al. Asgard archaea are the closest prokaryotic relatives of eukaryotes. PLoS Genetics forthcoming.

Spang A, et al. 2015. Complex archaea that bridge the gap between prokaryotes and eukaryotes. Nature 521: 173-+. doi: 10.1038/nature14447

Su X, et al. 2012a. YBR246W is required for the third step of diphthamide biosynthesis. J Am Chem Soc 134: 773–776. doi: 10.1021/ja208870a

Su X, et al. 2012b. Chemogenomic approach identified yeast YLR143W as diphthamide synthetase. Proc Natl Acad Sci U S A 109: 19983–19987. doi: 10.1073/pnas.1214346109

Suematsu T, et al. 2010. A bacterial elongation factor G homologue exclusively functions in ribosome recycling in the spirochaete Borrelia burgdorferi. Mol Microbiol 75: 1445–1454. doi: 10.1111/j.1365-2958.2010.07067.x

Suzek BE, et al. 2015. UniRef clusters: a comprehensive and scalable alternative for improving sequence similarity searches. Bioinformatics 31: 926–932. doi: 10.1093/bioinformatics/btu739

Tsuboi M, et al. 2009. EF-G2mt is an exclusive recycling factor in mammalian mitochondrial protein synthesis. Mol Cell 35: 502–510. doi: 10.1016/j.molcel.2009.06.028

Uthman S, et al. 2013. The amidation step of diphthamide biosynthesis in yeast requires DPH6, a gene identified through mining the DPH1-DPH5 interaction network. PLoS Genet 9: e1003334. doi: 10.1371/journal.pgen.1003334

Waterhouse AM, Procter JB, Martin DM, Clamp M, Barton GJ 2009. Jalview Version 2—a multiple sequence alignment editor and analysis workbench. Bioinformatics 25: 1189–1191. doi: 10.1093/bioinformatics/btp033

Webb TR, et al. 2008. Diphthamide modification of eEF2 requires a J-domain protein and is essential for normal development. J Cell Sci 121: 3140–3145. doi: 10.1242/jcs.035550

Williams TA, Foster PG, Nye TM, Cox CJ, Embley TM 2012. A congruent phylogenomic signal places eukaryotes within the Archaea. Proc Biol Sci 279: 4870–4879. doi: 10.1098/rspb.2012.1795

Williams TA, et al. 2017. Integrative modeling of gene and genome evolution roots the archaeal tree of life. Proc Natl Acad Sci U S A 114: E4602–E4611. doi: 10.1073/pnas.1618463114

Wu M, Scott AJ 2012. Phylogenomic analysis of bacterial and archaeal sequences with AMPHORA2. Bioinformatics 28: 1033–1034. doi: 10.1093/bioinformatics/bts079

Wu YW, Simmons BA, Singer SW 2016. MaxBin 2.0: an automated binning algorithm to recover genomes from multiple metagenomic datasets. Bioinformatics 32: 605–607. doi: 10.1093/bioinformatics/btv638

Yang J, et al. 2015. The I-TASSER Suite: protein structure and function prediction. Nat Methods 12: 7–8. doi: 10.1038/nmeth.3213

Yu YR, You LR, Yan YT, Chen CM 2014. Role of OVCA1/DPH1 in craniofacial abnormalities of Miller-Dieker syndrome. Hum Mol Genet 23: 5579–5596. doi: 10.1093/hmg/ddu273

Zaremba-Niedzwiedzka K, et al. 2017. Asgard archaea illuminate the origin of eukaryotic cellular complexity. Nature 541: 353–358. doi: 10.1038/nature21031

Zhang Y, Liu S, Lajoie G, Merrill AR 2008. The role of the diphthamide-containing loop within eukaryotic elongation factor 2 in ADP-ribosylation by Pseudomonas aeruginosa exotoxin A. Biochem J 413: 163–174. doi: 10.1042/BJ20071083

